# Early development of the mineralized external skeleton of the polyplacophoran mollusk, with insight into the evolutionary history of shell plates and spicules

**DOI:** 10.1101/2024.05.20.594941

**Authors:** Hiroki Yoshikawa, Yoshiaki Morino, Hiroshi Wada

## Abstract

Recent molecular phylogenetic studies have raised two questions about the evolutionary history of the calcified exoskeleton of mollusks. The first question concerns the homology of the two types of skeleton; whether spicules and shell plates share an evolutionary origin. The second question is the homology of the shell plates between chitons and other mollusks, including gastropods and bivalves. To gain insight into these questions, we examined the early development of shell plates and spicules in chitons. We identified several developmental genes that are involved in both shell plates and spicules, suggesting that spicules and shell plates share a common evolutionary origin. We also found that subpopulations of the dorsal shell field (the ridge and the plate field) have specific gene expression profiles. The differential gene expression of the ridge and plate field is not identical to the profiles of the zones of the gastropod shell field. This observation may suggest an independent evolutionary origin of the shell plates in chitons and gastropods.

## Introduction

The Cambrian explosion is one of the most dramatic events in the evolutionary history of life, often thought to have been facilitated by an arm race between predators and defenders. As part of the defence system, multicellular animals independently acquired biomineralised skeletal structures in several lineages, such as arthropods, bryozoans, echinoderms and vertebrates. Mollusks are also a representative group of animals that acquired skeletal structures, diverging by the acquisition and further elaboration of the calcified exoskeleton.

Although mollusks are often thought of as animals with a shell plate(s), some mollusks lack a shell plate, such as Aplacoholans: Caudofoveata and Solenogastres. Instead, their body surface is covered by spicules, which also consist of calcium carbonate (Todt et al., 2008). Aplacophorans were once considered a basal group of mollusks. This phylogenetic framework favoured the idea that spicules preceded the acquisition of shell plates and that shell plates evolved as an elaboration of spicules (Haszprunar, 2000). However, some authors did not support the evolutionary affinity between spicules and shell plates, and rather suggested independent origins of two mineralised tissues (Scheltema and Schander, 2006; Kocot, 2013).

In the argument for the evolution of mineralised tissues in molluscs, the chiton occupies a unique position because it possesses both shell plates and spicules. According to a recent molecular phylogenetic analysis (Kocot et al., 2011), molluscs are divided into two main groups: Conchifera and Aculifera. The former includes gastropods, bivalves, scaphopods, monoplacophorans and cephalopods, while the latter includes aplacophorans and polyplacophorans. The Conciferans are characterized by the presence of shell plate(s), while the latter are characterized by the presence of spicules. This phylogeny revived the argument of the evolutionary history of molluscan mineralized tissues, because the phylogenetic framework supported the hypothesis that spicules were acquired in the lineage towards Aculifera, and that it is equally parsimonious to consider the single origin of shell plates in the common ancestors of molluscs (with a single loss of it in aplacopholans), or the independent acquisition of shell plates in chiton and conciferans (Kocot et al., 2011). In order to elucidate the evolutionary history of shell plates, a more comprehensive comparison of the developmental mechanisms of shell plates and spicules is therefore required. For the analysis, the chiton should provide critical knowledge, because they develop both structures from the single set of genomes.

The Conciferan shell is covered by an organic layer, the periostracum, which is secreted from the outer zone of the mantle edge, the periostracal groove (Kocot et al., 2016). The underlying shell matrix is secreted by the epithermal cells of the inner zone of the mantle margin. Similar zonation is also observed in the larval shell field, although the clear functional differentiation of the zones has yet to be elucidated (Johnson et al., 2019). Shell plates and spicules of the chiton are also covered by an organic layer. (Kniprath, 1980) reported the zonation of the shell field of the chiton larva, and the cells in the ridge of the shell field were suggested to be responsible for the secretion of the organic layer (cuticle). The shell matrix was suggested to be secreted by the cells of the interridge, referred to as the plate field.’’ Checa et al. (2017) identified two cell types in the girdle field of chiton where spicules develop. The columnar cells of the papillae were proposed to be involved in the secretion of the spicule matrix, while the cuboidal cells of the girdle epithelium secrete the organic layer. Although the organic layer plays a critical role in the formation of the skeletal matrix, the homology of the organic layers between chiton and conciferans is controversial (Checa et al., 2017).

Previous studies have suggested that chiton shell development is regulated by the use of genes involved in conciferan shells, such as *engrailed, hox* genes and *Pax2/5/8* (Huan et al., 2020; Jacobs et al., 2000; Wollesen et al., 2017; Xia et al., 2023). Although these studies support the homology of shell plate(s) between Chiton and Conciferans, little is known about the developmental mechanisms of spicules. In this study, we examined the expression of several genes whose conciferan homologs mark their shell field. Our study showed that the majority of genes involved in the development of the shell field are also expressed in the girdle field and are therefore likely to be involved in the development of spicules in chiton. This observation supports the evolutionary affinity between spicules and shell plates. We also found that in the shell field, the girdle and plate fields show a distinct set of gene expression, supporting the previous report of functional differentiation between the girdle and plate fields. Interestingly, we found that the gene set for the ridge or plate field does not correspond to the set of genes that show different expression patterns in the zones of the conciferan shell field. Evolutionary history of skeletal structures is discussed based on gene expression profiles.

## Results and Discussion

The development of the chiton, *Acanthochitona* sp.A., is similar to that described for *A. rublolineata* (Xia et al., 2023) (Fig. 1). The first signs of spicule development were observed on the dorsal side, just anterior to the prototroch, as well as in the posterior end of the larva at 28 hpf (Fig.1G). At 48 hpf, birefringent spicules were observed in the dorsal side of the pretotrochal region and lateral side of the posttrochal epidermis, described as the outer girdle field in (Xia et al., 2023) (Fig.1H). The matrix of shell plates was not clearly observed at this stage. Metamorphosis is complete before 144 hpf and seven shell plates are clearly observed in the dorsal epidermis (Fig.1I).

**Figure 1.**
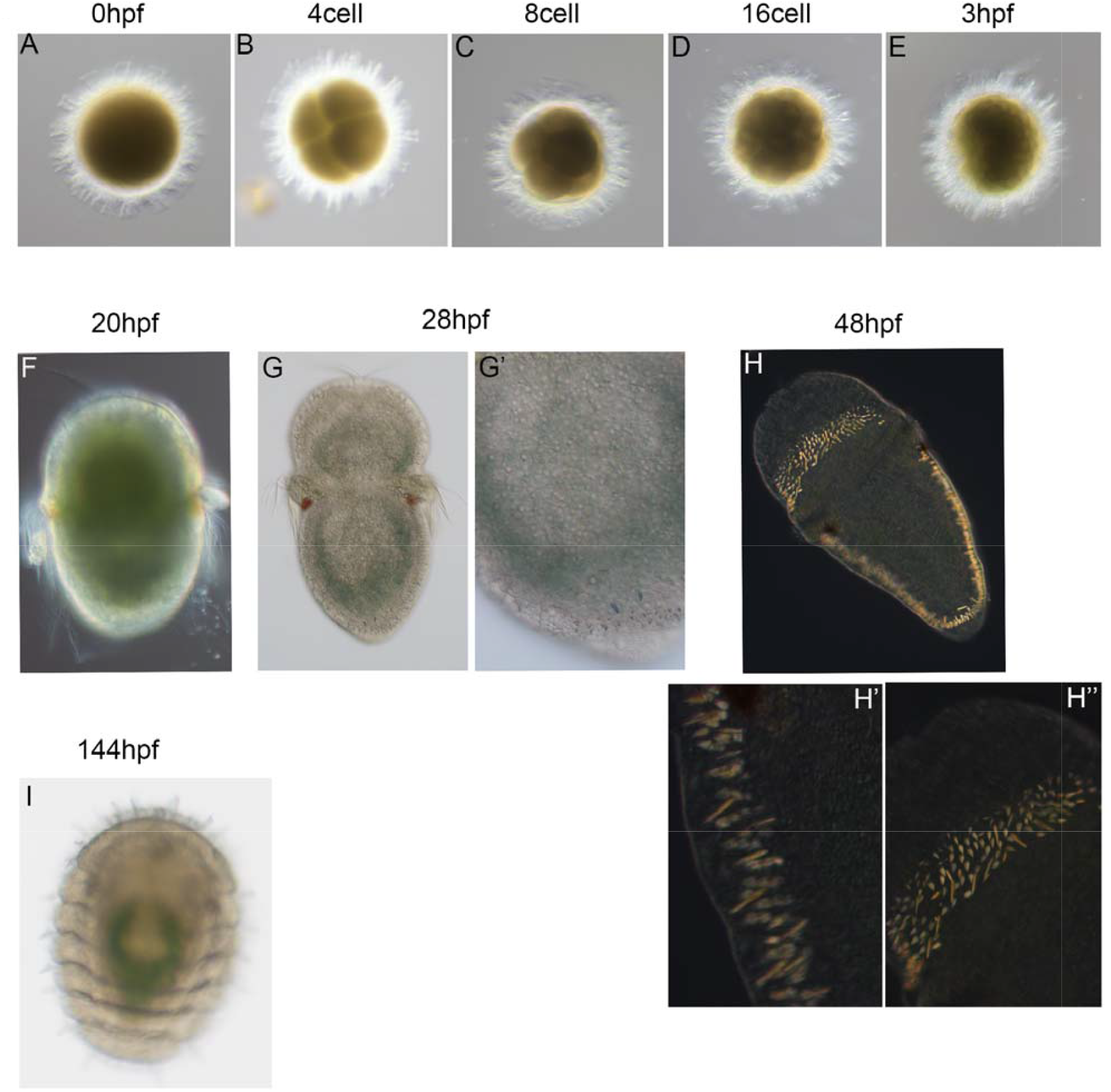
Developmental processes in *A*.sp.A. (A-E) The early cleavage stages. (F) The newly hatched larvae of 20 hpf. No shell plate or sclerite are visible at this stage. (G) 28 hpf larva show small spicules in the posterior end. (H) Spicules are visible in pretrochal regiona as well as in the lateral girdle region of 48 hpf larvae. (I) Juvenile of 144 hpf.

### Shell matrix protein (SMP) in the shell field and the girdle field in trochophore larvae

To identify the cells involved in chiton skeletogenesis, we first examined the expression of genes whose homologs are known to encode shell matrix proteins. We identified *chitin synthase* (*cs*) and *Pif* (Suzuki et al., 2009) from the transcriptome data of *Acanthochitona* sp. A (Morino and Yoshikawa, 2023). We found that both are expressed in the cells of seven stripes of the shell field in the posttrochal region of 45 hpf larvae (Figure 2A-F). Close observation of the expression revealed that both are expressed in the ridge of the shell field (Fig. 2C, F). In addition, *cs* and *Pif* are also expressed in the pretrochal region and in the outer girdle region of 45 hpf larvae, where spicules develop (Fig. 2B, E).

**Figure 2.**
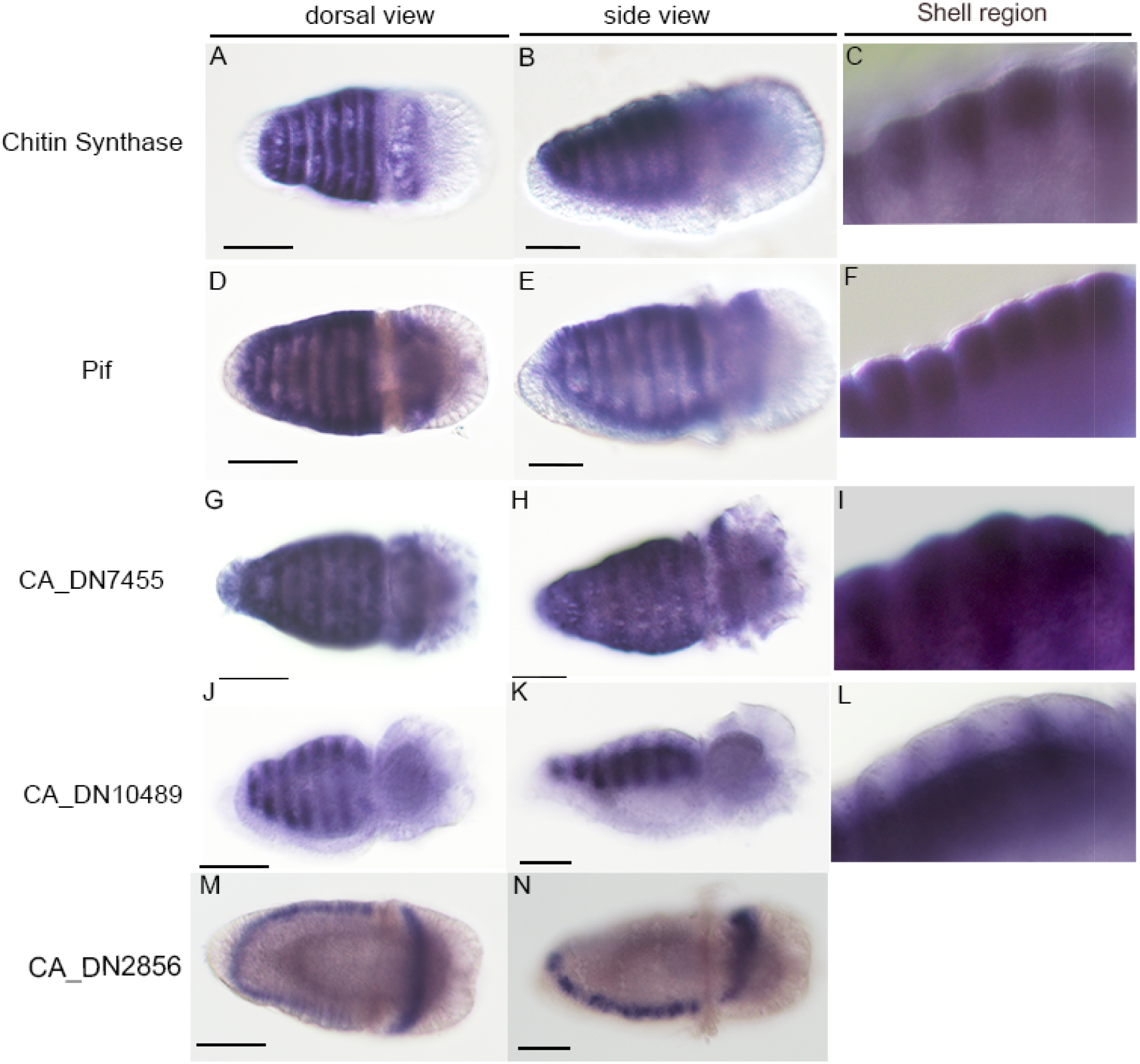
The expression patterns of the shell matrix proteins (SMPs) The expression patterns of the shell matrix proteins in the dorsal view and the side view. Magnified images were shown in C, F, I, L. Scale bars: 50 μm.

From the transcriptome data of trochophore larvae (Morino and Yoshikawa, 2023), we identified three carbonic anhydrase (CA) genes. *CA7455* shows a similar expression pattern to *cs* and *Pif* in the ridge of the shell field, the pretrochlear epidermis and the outer girdle field (Fig. 2G-I). *CA10489* is expressed in the plate field (inter-ridge) of the shell field (Xia et al., 2023), and no expression is detected in either the pretrochal region or the girdle field (Figs. 2J-L). In contrast, *CA2856* is expressed in the pretrochal epidermis and the outer girdle field, but not in the shell field (Fig. 2M, N).

Kniprath (1980) identified four types of cells in the chiton shell field, type 1-2 in the ridge, and type 3-4 in the inter-ridge plate field. The former were proposed to be responsible for the secretion of the organic layer and the latter for the secretion of the shell matrix. This does not agree well with our observation that *cs* is expressed in the ridge, because Furuhashi et al. (2009) reported that chitin is not contained in the organic layer of the chiton shell. The expression of *pif* in the ridge is more consistent with the idea that the ridge of the larval chiton is responsible for the shell matrix rather than the matrix for the organic layer.

### Differential expressions of transcription factors in the shell field and the girdle field of chiton

We then examined the expression of genes encoding transcription factors whose orthologs are expressed in the shell field of gastropods or bivalves: *Gbx, Grainyhead, engrailed, Gata1/2/3, Pax2/5/8, Hox1* and *Goosecoid*. We found that all seven transcription factor genes are expressed in the shell field of 45 hpf larvae (Fig. 3). While *Hox1* shows stronger expression in the anterior stripes, the rest of the genes show uniform expression in seven stripes (Fig. 3). Closer examination of the expression revealed that *Gbx* and *Grainyhead* are expressed in the plate field cells (Fig. 3C, L). The other five genes show expression in the ridge region of the plate field (Fig. 3F, I, C, R). *engrailed* expression in the ridge of the plate field has been reported in *A. rubrolineata* (Xia et al., 2023). Pretrochal expression and outer girdle field expression are observed for *Grainyhead, Gata1/2/3, Pax2/5/8, Goosecoid. Hox1* does not show any expression in the pretrochlear region (Fig. 3). The girdle field expression of *Hox1* is stronger in the anterior region (Fig. 3P, Q). For *Gbx* and *engrailed*, expression was not detected in either the pretrochal region or the girdle field.

**Figure 3.**
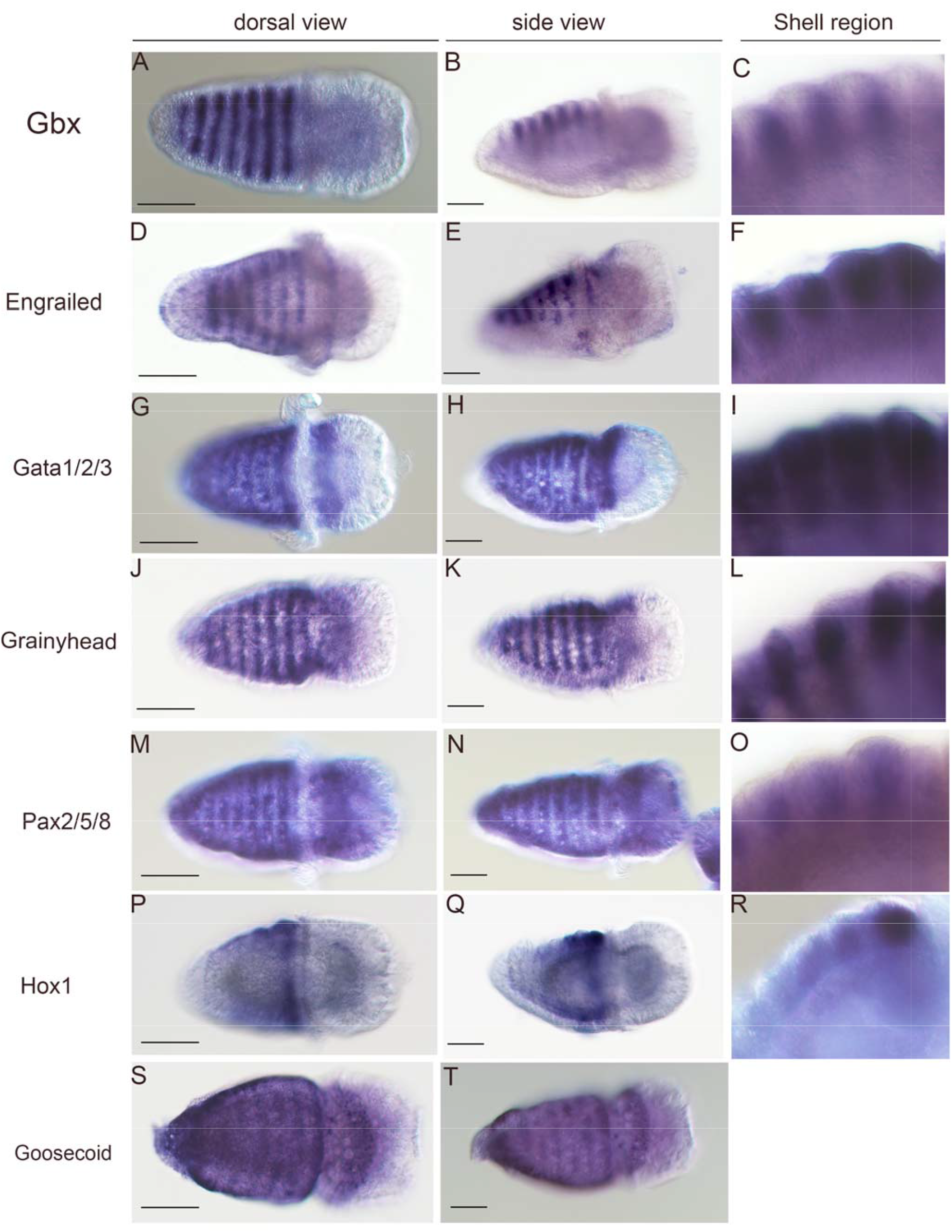
The expression patterns of the transcription factor (TFs) The expression patterns of the transcription factor genes in the dorsal view and the side view. Magnified images were shown in C, F, I, L, O, R. Scale bars: 50 μm.

### Evolutionary Implications for the molluscan external skeletons

The above results show that five transcription factor genes are expressed both in the shell field and in the regions of spiculogenesis (the pretrochal epidermis and the outer girdle field). This supports the evolutionary affinity between the two types of chiton skeleton, the shell plates and the spicules. Together with *engrailed*, six transcription factor genes are also expressed in the conciferan shell field. Therefore, the development of molluscan skeletons is regulated by similar repertoires of transcription factors, suggesting a single origin of molluscan skeletogenesis.

However, this does not necessarily support a single origin of the shell plates in conciferans and chitons. Rather, we find that two zones of the shell field are not specified by the same set of transcription factors in conciferans and chiton. We showed that *Gbx* and *Grainyhead* specify the plate field and the remaining five genes specify the ridge. Our single cell transcriptome analysis of the gastropod *Nipponacmea* showed that *Hox1* and *engrailed* mark the inner domain of the gastropod shell field, while *Gata1/2/3, Pax2/5/8* and *Goosecoid* mark the outer domain of the shell field (Phuangphong et al., 2024). Johnson et al. (2019) also shows that *Hox1* and *Pax2/5/8*-*Goosecoid* mark different zones of the shell field in the other species of gastropod *Tritia*. These observations may be more consistent with the idea that shell field zonation occurred independently in chiton and conciferans, and in turn support the independent evolutionary origin of shell plates between chitons and conciferans.

## Materials and Methods

### 1. Animal Collection and Cultures

Adult specimen of chiton *Acanthochitona* sp. A was collected from the coastal area of Hiraiso (Ibraki, Japan) during the breeding season (June - July). Details of species identification of *Acanthochitona* was described in (Morino and Yoshikawa, 2023). Fertilized egges were obtained from sperms and eggs spontaneously spawned several hours after collection. Embryos were reared in artificial seawater at 25 °C.

### 2. Gene survey and Phylogenetic analysis

Genes encoding the transcription factors and Pif were identified by Blast (BLAST 2.13.0+) search against the transcriptome data reported in (Morino and Yoshikawa, 2023). CA and chitin synthase sequences were extracted by HMMER search (ver3.4) using PF03142.16 (Chitin_synth_2) and PF00194.22 (Carb_anhydrase) against the translated transcriptome (Morino and Yoshikawa, 2023).

The orthology of extracted genes of *A*. sp.A was confirmed by the phylogenetic analysis by the method described previously (Morino and Yoshikawa, 2023)(Fig. S1-S7).

### 3. Whole mount in situ hybridization

cDNAs for probes were amplified by using primers listed in Table S1. Whole-mount in situ hybridization was performed using the method described previously (Morino and Yoshikawa, 2023), with the ProK concentration modified to 60 ug/ml.

## References

Checa, A.G., Vendrasco, M.J., Salas, C., 2017. Cuticle of Polyplacophola: structure, secretion, and homology with the periostracum of conciferans. Mar. Biol. 164, 64.

Furuhashi, T., Schwarzinger, C., Miksik, I., Smrz, M., Beran, A., 2009. Molluscan shell evolution with review of shell calcification hypothesis. Comp. Biochem. Physiol. B 154, 351–371.

Haszprunar, G., 2000. Is the Aplacophora monophyletic? A cladistic point of view. Am. Malacol. Bull. 15, 115–130.

Huan, P., Wang, Q., Tan, S., Liu, B., 2020. Dorsoventral decoupling of Hox gene expression underpins the diversification of molluscs. Proc. Natl. Acad. Sci. USA 117, 503–512.

Jacobs, D.K., Wray, C.G., Wedeen, C.G., Kostriken, R., DeSalle, R., Staton, J.L., Gates, R.D., Lindberg, D.R., 2000. Molluscan engrailed expression, serial organization, and shell evolution. Evol. Dev. 2, 340–347.

Johnson, A.B., Fogel, N.S., Lambert, J.D., 2019. Growth and morphogenesis of the gastropod shell. Proc. Natl. Acad. Sci. USA 116, 6878–9883.

Kniprath, E., 1980. Ontogenetic plate and plate field development in two chitons, Middendorffia and Ischnochiton. Roux’s Archiv. Dev. Biol. 189, 97–106.

Kocot, K.M., 2013. Recent Advances and unanswered questions in deep molluscan phylogenetics. Am. Malacol. Bull. 31, 1–14.

Kocot, K.M., Aguilera, F., McDougall, C., Jackson, D.J., Degnan, B.M., 2016. Sea shell diversity and rapidly evolving secretomes: insights into the evolution of biomineralization. Front. Zool. 13, 23.

Kocot, K.M., Cannon, J.T., Todt, C., Citarella, M.R., Kohn, A.B., Meyer, A., Santos, S.R., Schander, C., Moroz, L.L., Lieb, B., Halanych, K.M., 2011. Phylogenomics reveals deep molluscan relationships. Nature 477, 452–456.

Morino, Y., Yoshikawa, H., 2023. Role of maternal spiralian-specific homeobox gene SPILE-Ein the specification of blastomeres along the animal–vegetalaxis during the early cleavage stages of mollusks. Dev. Growth Differ. 65, 384–394.

Phuangphong, S., Yoshikawa, H., Kojima, Y., Wada, H., Morino, Y., 2024. Development of shell field populations in gastropods. biorxiv 10.1101/2024.05.16.592602.

Scheltema, A.H., Schander, C., 2006. Exokeletons: tracinf molluscan evolution. Venus 65, 19–26.

Suzuki, M., Saruwatari, K., Kogure, T., Yamamoto, Y., Nishimura, T., Kato, T., Nagasawa, H., 2009. An Acidic Matrix Protein, Pif, Is a Key Macromolecule for Nacre Formation. Science 325, 1388–1390.

Todt, C., Okusu, A., Schander, C., Schwabe, E., 2008. Solenogastres, Caudofoveata and Polyplacophola, in: Ponder, W.F., Lindberg, D.R. (Eds.), Phylogeny and Evolution of the Mollusca. University od California Press, Berkeley.

Wollesen, T., Scherholz, M., Monje, S.V.R., Redl, E., Todt, C., Wanninger, A., 2017. Brain regionalization genes are co-opted into shell field patterning in Mollusca. Sci. rep. 7, 5486.

Xia, Y., Huan, P., Liu, B., 2023. Shell field morphogenesis in the polyplacophoran mollusk Acanthochitona rubrolineata. EvoDevo 14, 5.

