## Supplement for "Early development of the mineralized external skeleton of the polyplacophoran mollusk, with insight into the evolutionary history of shell plates and spicules"

**List of Supplementary Materials**

**Figure S1**

Molecular phylogenetic tree of Hox genes

**Figure S2**

Molecular phylogenetic tree of Engrailed and Gbx, Dlx genes

**Figure S3**

Molecular phylogenetic tree of Gata genes

**Figure S4**

Molecular phylogenetic tree of Pax genes

**Figure S5**

Molecular phylogenetic tree of Grainyhead and LSF genes

**Figure S6**

Molecular phylogenetic tree of Goosecoid and Otx genes

**Figure S7**

Molecular phylogenetic tree of Pif-like genes

**Table S1**

Primer used for gene isolation

**Table S2**

List of chitin synthase and Carbon anhydrase and their expression levels (TPM) in A sp.A

**DatasetS1.fasta**

Aligned sequences of transcription factor and Pif-like for phylogenetic analysis

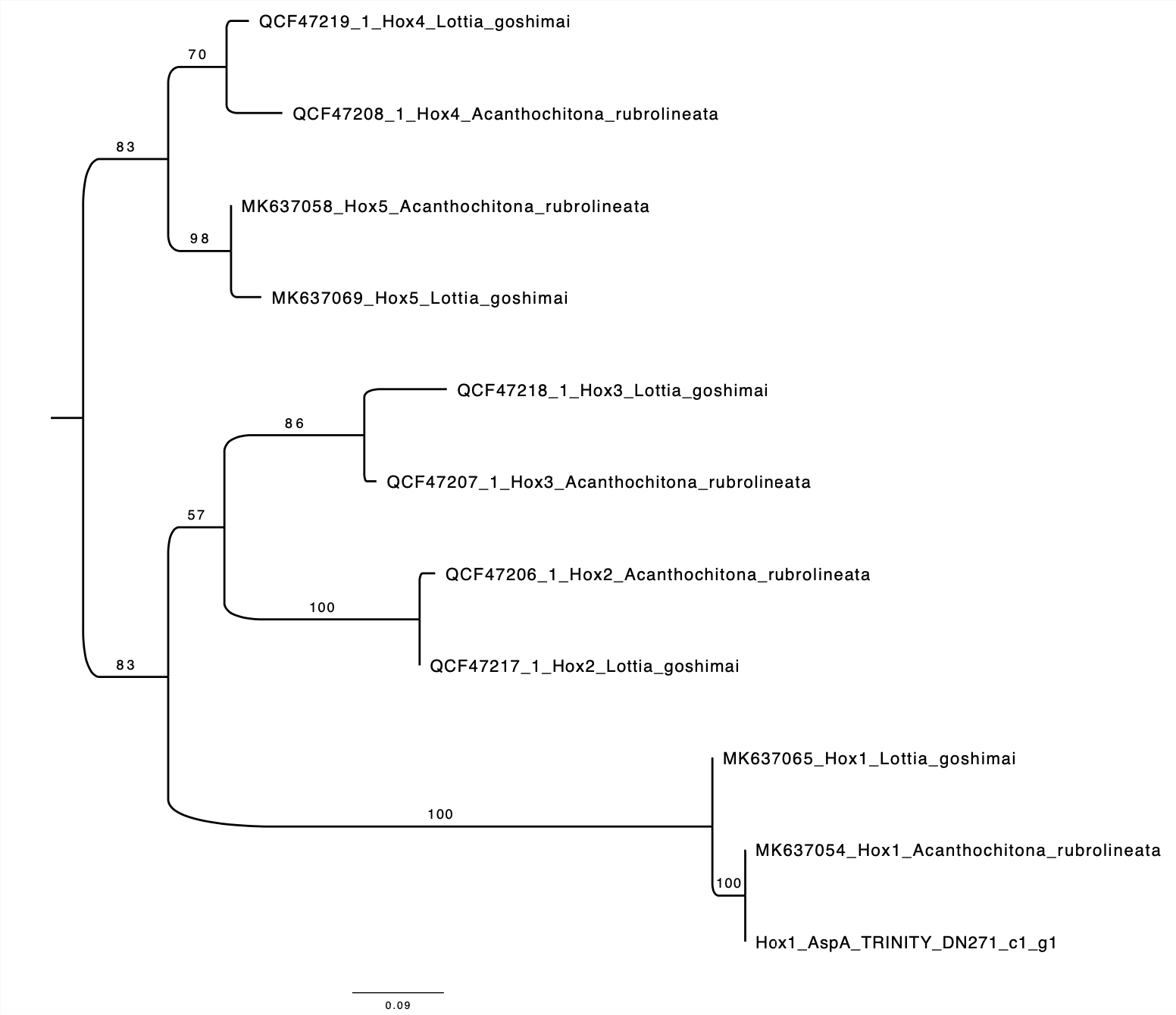

**Fig. S1 Molecular phylogenetic tree of Hox genes**

A molecular phylogenetic tree of the Hox gene of *Acanthochitona* sp. A and previously annotated Hox genes of mollusks (1) (Huan et al. (2019)) was constructed. The amino acid sequences of the homeodomains were used to construct a tree based on the maximum likelihood method. LG was selected by RaxML and used as an amino acid substitution model. The numbers at the nodes are the bootstrap values from 100 replicates. The trees were visualized by FigTree (http://tree.bio.ed.ac.uk/software/figtree/).

AspA: *Acanthochitona* sp. A

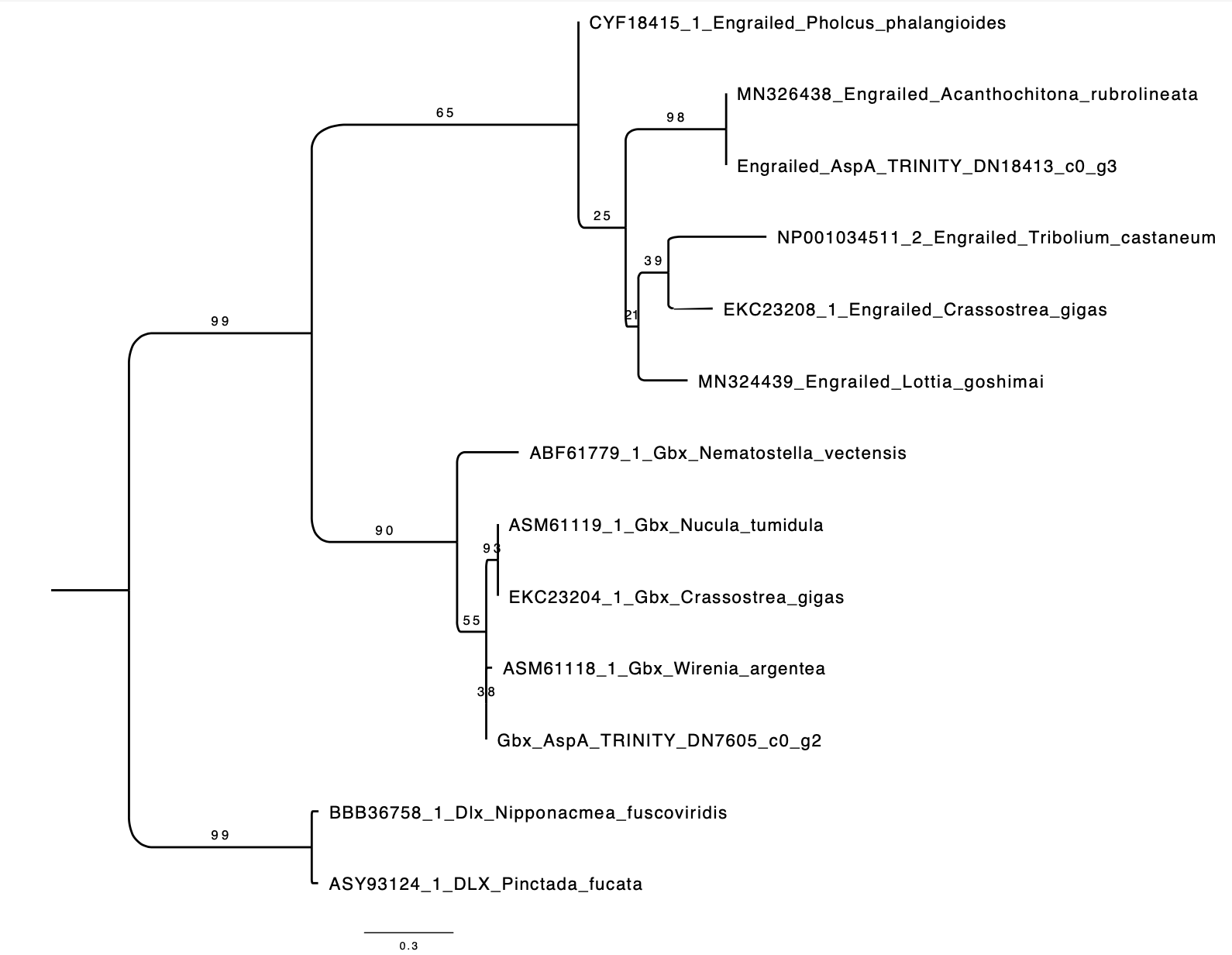

**Figure S2 Molecular phylogenetic tree of Engrailed and Gbx, Dlx genes**

A molecular phylogenetic tree of the Engrailed and Gbx gene of *Acanthochitona* sp. A. We used Dlx genes for outgroups. The amino acid sequences of the homeodomains were used to construct a tree based on the maximum likelihood method. LG was selected by RaxML and used as an amino acid substitution model. The numbers at the nodes are the bootstrap values from 100 replicates. The trees were visualized by FigTree (http://tree.bio.ed.ac.uk/software/figtree/).

AspA: *Acanthochitona* sp. A

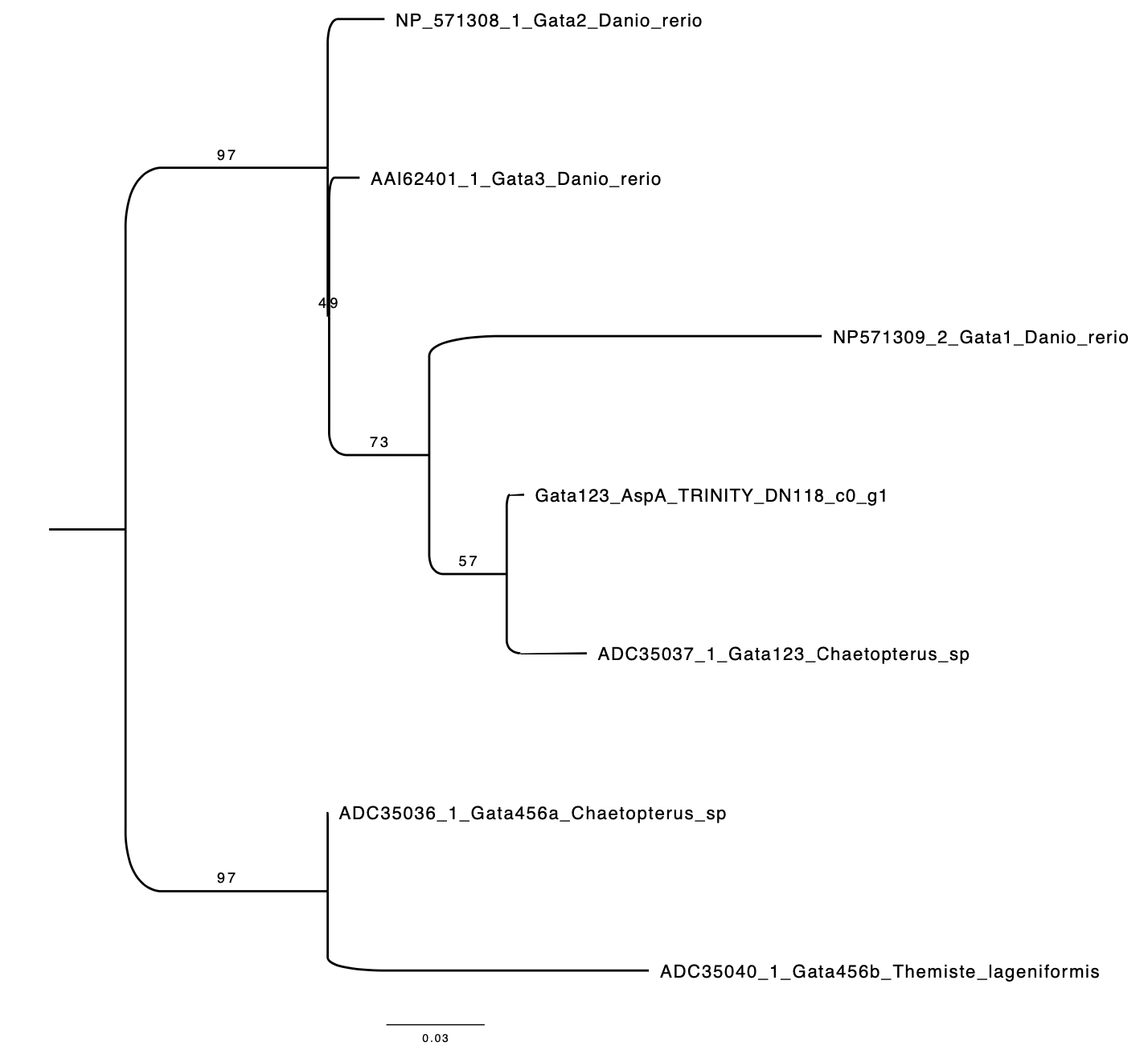

**Figure S3 Molecular phylogenetic tree of Gata genes**

A molecular phylogenetic tree of the Gata1/2/3 gene of *Acanthochitona* sp. A. We used Gata4/5/6 genes for outgroups. The amino acid sequences of the homeodomains were used to construct a tree based on the maximum likelihood method. LG was selected by RaxML and used as an amino acid substitution model. The numbers at the nodes are the bootstrap values from 100 replicates. The trees were visualized by FigTree (http://tree.bio.ed.ac.uk/software/figtree/).

AspA: *Acanthochitona* sp. A

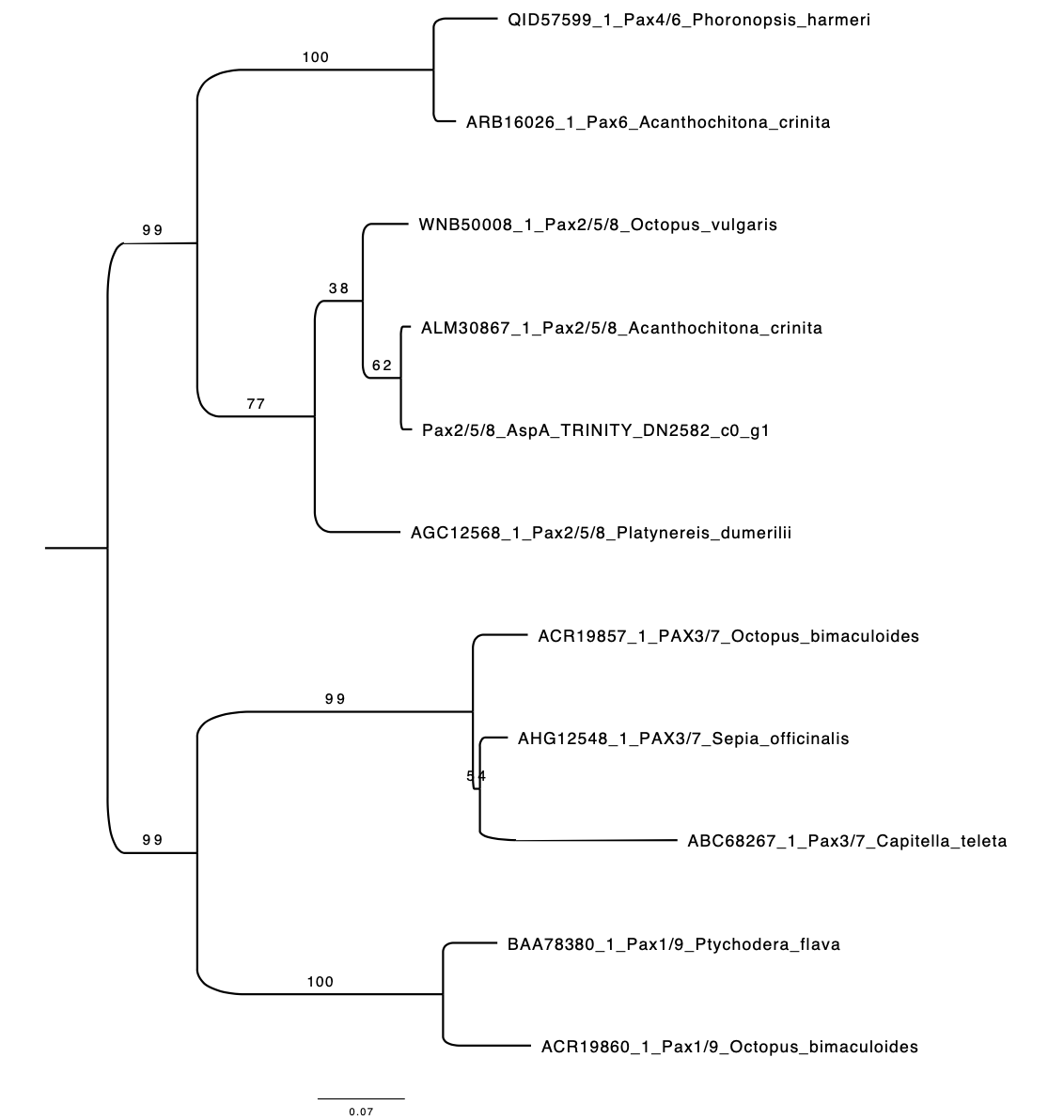

**Figure S4 Molecular phylogenetic tree of Pax genes**

A molecular phylogenetic tree of the Pax gene of *Acanthochitona* sp.A. The amino acid sequences of the paired domain were used to construct a tree based on the maximum likelihood method. LG was selected by RaxML and used as an amino acid substitution model. The numbers at the nodes are the bootstrap values from 100 replicates. The trees were visualized by FigTree (http://tree.bio.ed.ac.uk/software/figtree/).

AspA: *Acanthochitona* sp. A

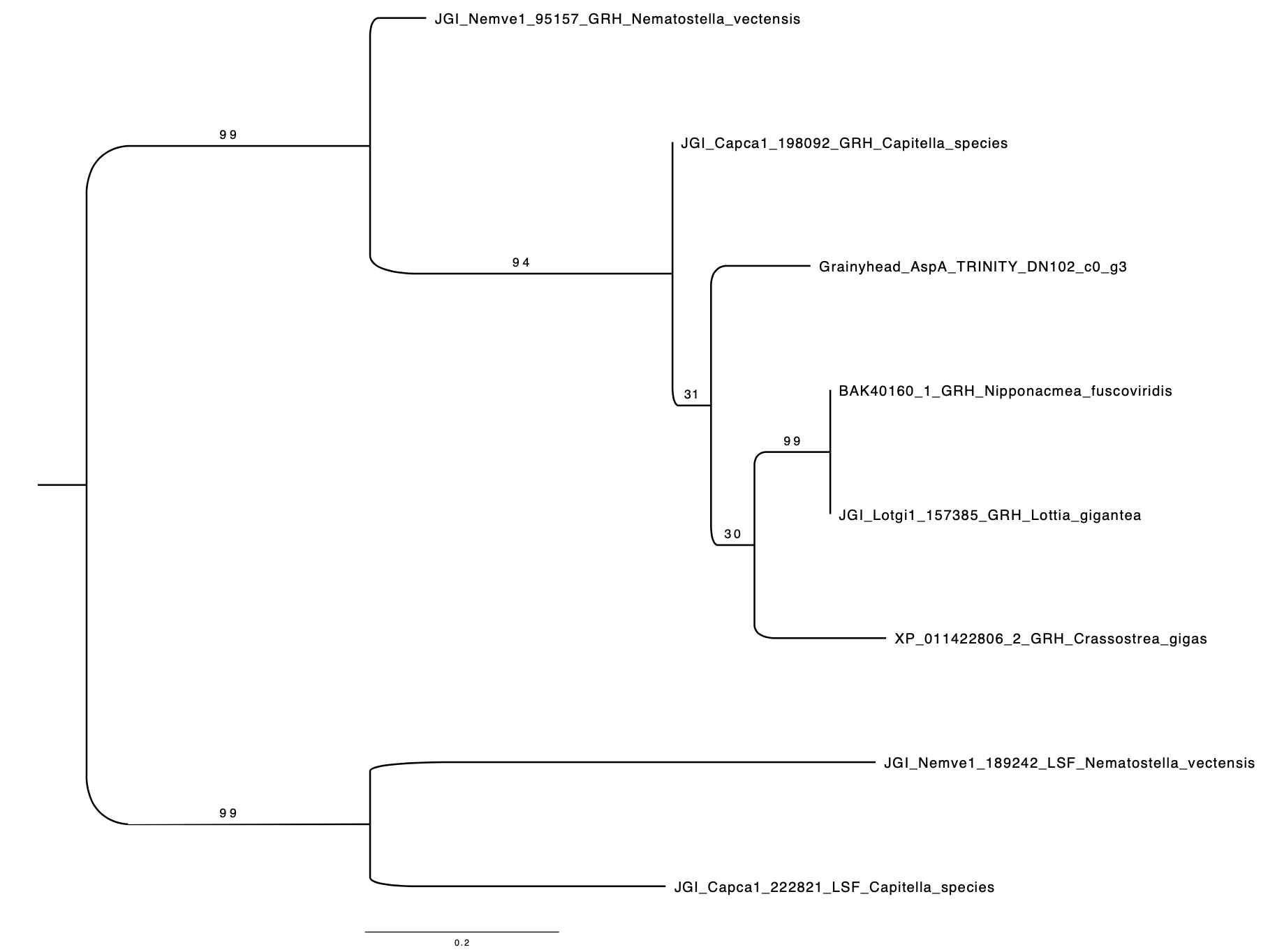

**Figure S5 Molecular phylogenetic tree of Grainyhead and LSF genes**

A molecular phylogenetic tree of the grainyhead(GRH) gene of *Acanthochitona* sp.A. We used LSF genes for outgroups(2) (Traylor-Knowles et al., 2010). The amino acid sequences of the paired domain were used to construct a tree based on the maximum likelihood method. LG was selected by RaxML and used as an amino acid substitution model. The numbers at the nodes are the bootstrap values from 100 replicates. The trees were visualized by FigTree (http://tree.bio.ed.ac.uk/software/figtree/).

AspA: *Acanthochitona* sp. A

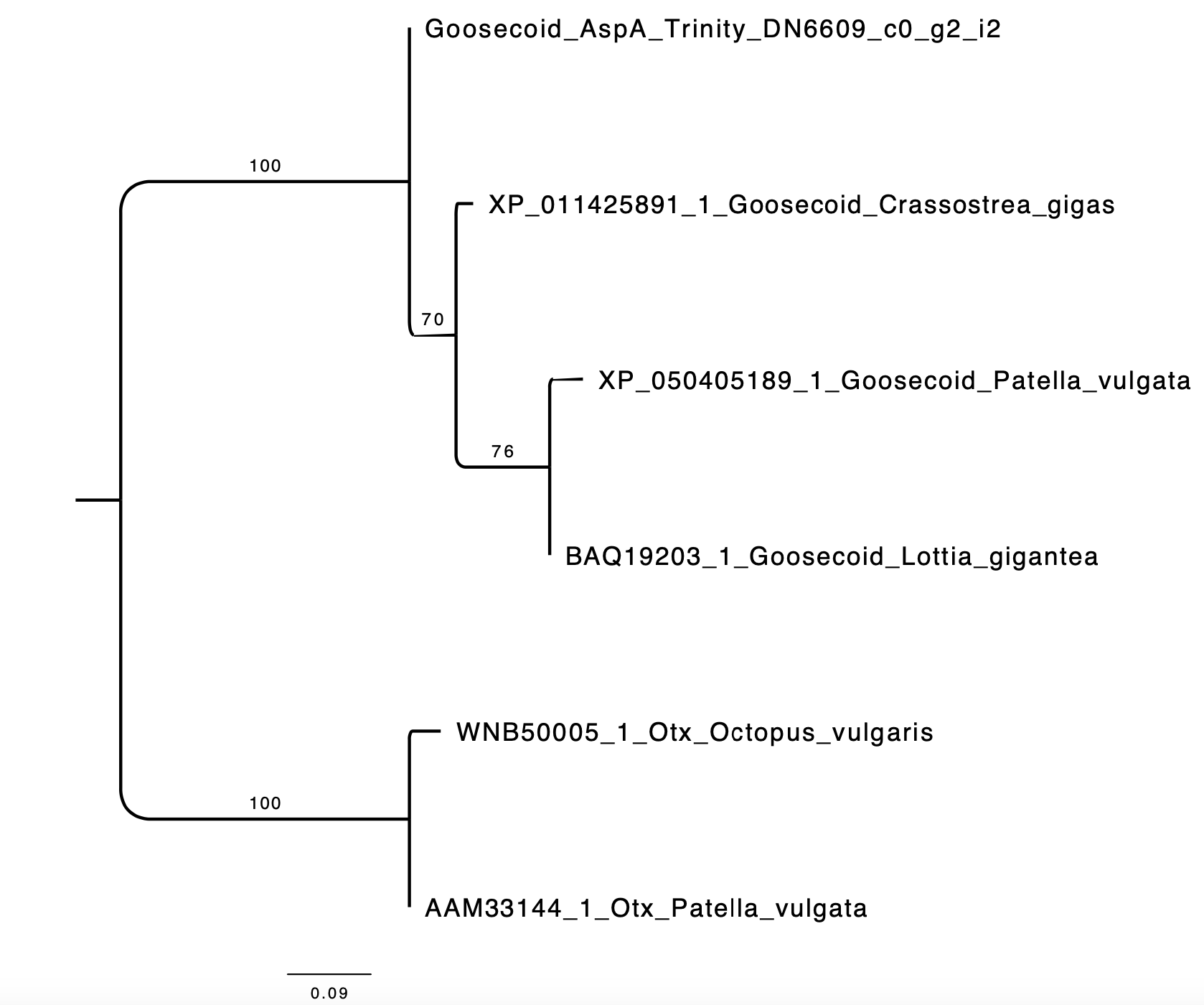

**Figure S6 Molecular phylogenetic tree of Goosecoid and Otx genes**

A molecular phylogenetic tree of the Goosecoid gene of *Acanthochitona* sp.A. We used Otx genes for outgroups. The amino acid sequences of the paired domain were used to construct a tree based on the maximum likelihood method. LG was selected by RaxML and used as an amino acid substitution model. The numbers at the nodes are the bootstrap values from 100 replicates. The trees were visualized by FigTree (<http://tree.bio.ed.ac.uk/software/figtree/>).

AspA: *Acanthochitona* sp. A

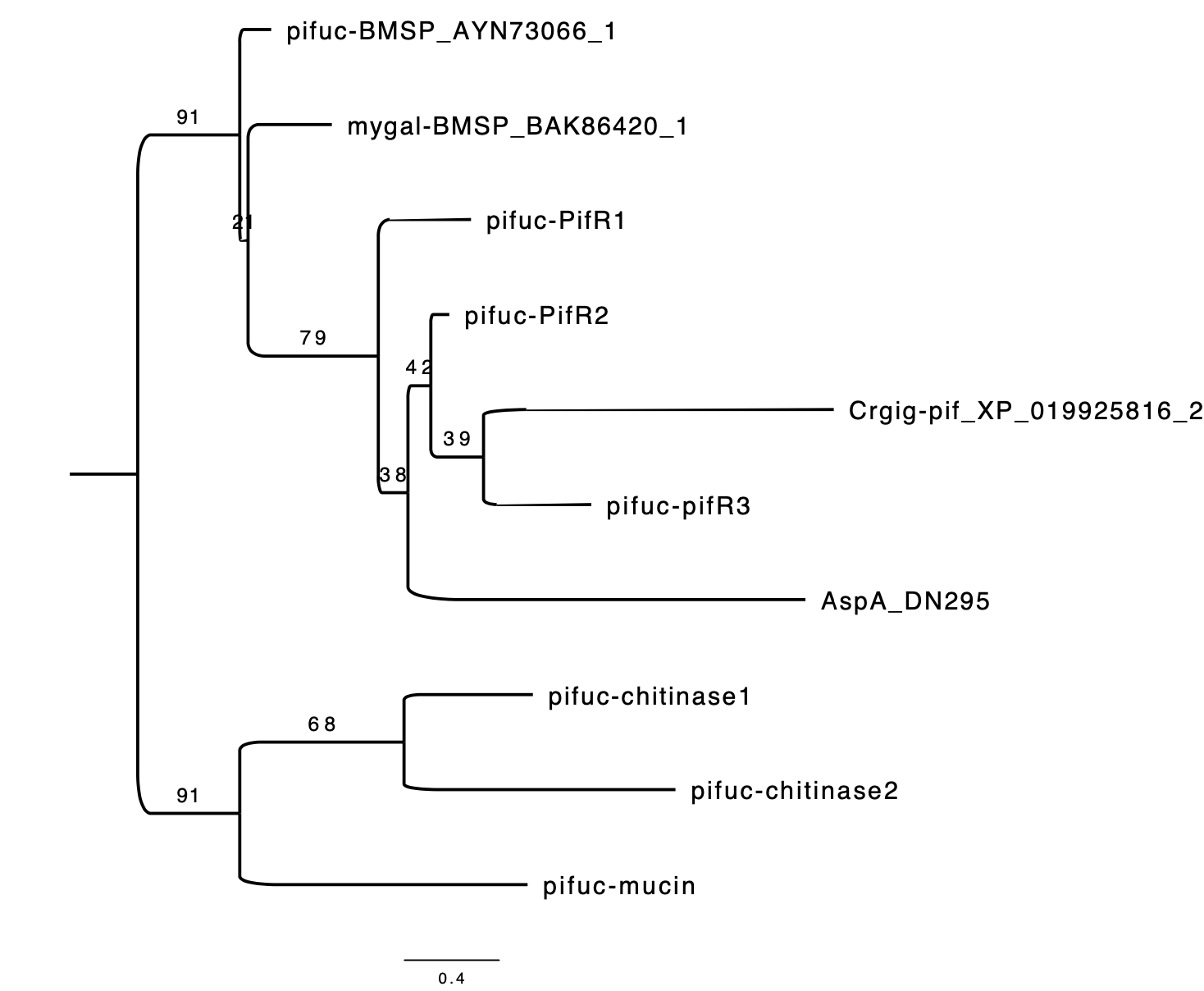

**Figure S7 Molecular phylogenetic tree of Pif-like genes**

A molecular phylogenetic tree of the Pif-like gene of *Acanthochitona* sp.A. Phylogenetic analysis of Pif was performed using aligned and used genes, as in previous studies (3)(Miyamoto et al., 2013). The amino acid sequences of the paired domain were used to construct a tree based on the maximum likelihood method. LG was selected by RaxML and used as an amino acid substitution model. The numbers at the nodes are the bootstrap values from 100 replicates. The trees were visualized by FigTree (http://tree.bio.ed.ac.uk/software/figtree/).

Pifuc: *Pinctada fucata*; Crgig: *Crassostrea* *gigas*; mygal: *Mytilus galloprovincialis*

AspA: *Acanthochitona* sp. A

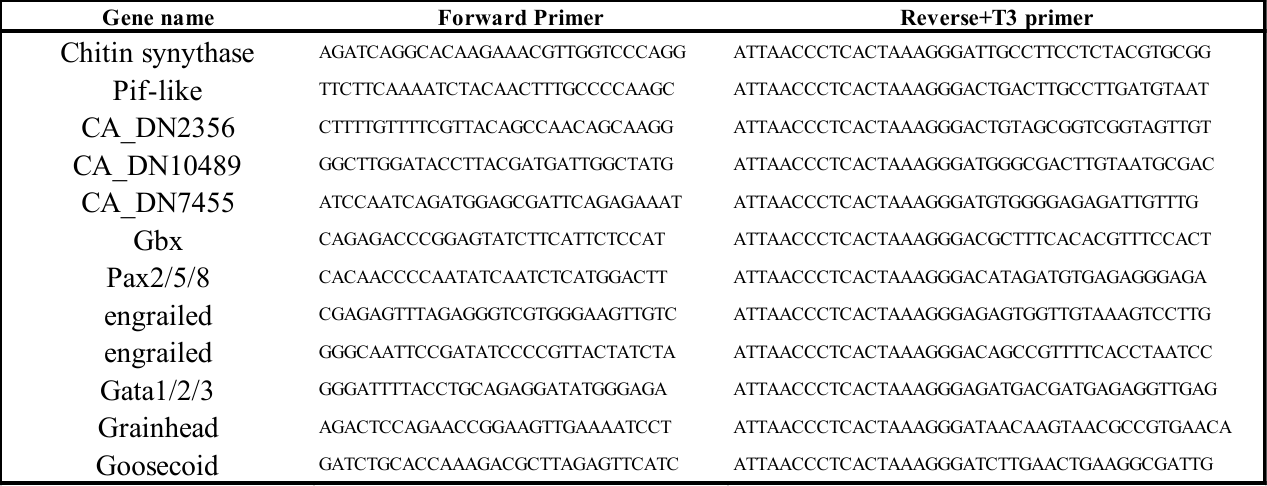

**Table S1 Primer used for gene isolation**

**References**

1.	P. Huan, Q Wang, B Liu, Dorsoventral decoupling of Hox gene expression underpins the diversification of molluscs. *PANS***117**,503-512, (2019).

2.	N. Traylor-Knowles, U. Hansen, T. Q. Dubuc, L. Kaufman, J. R. Finnerty, The evolutionary diversification of LSF and Grainyhead transcription factors preceded the radiation of basal animal lineages. *BMC Evol Biol* **10**, 101 (2010).

3.	H. Miyamoto *et al*., The diversity of shell matrix proteins: Genome-wide investigation of the pearl oyster, *Pinctada fucata*. *Zool Sci* **30**, 801-816 (2013).
